# Intralimb locomotor coordination in rats walking on asymmetric pegway

**DOI:** 10.1101/2023.01.27.525960

**Authors:** Kacie Hanna, Ezequiel M Salido, Neha Lal, Kiril Tuntevski, Sergiy Yakovenko

## Abstract

Complex movements such as walking or reaching are generated by a sequence of muscle actions. How these coordinated actions subserve complex movements and their recovery after disruption remains unknown. The use of high throughput recording-stimulation systems with microelectrode access to structures along the neuraxis may complement the neurological models in rodents. To this purpose, we have trained rats to perform the precise foot placement locomotor task that allows us to assess skilled locomotor movements. Animals were pretrained on the peg walkway task, which was configured to impose either symmetric or asymmetric (with overstepping) locomotor stepping at preferred stride length. Selected forelimb muscles were implanted with intramuscular differential electrodes. After a week of recovery, we collected electromyography from the implanted muscles and ground reaction forces from the array of force sensors embedded into walkway pegs. The temporal relationship between muscle bursts was measured for each intralimb set of muscles (n=13) in symmetric and asymmetric stepping. The sequence corresponded to the progression of muscle actions responsible for limb lift, flexion and transport, overground clearance, and preparation for ground contact. The stereotyped spatiotemporal sequence of muscle activity was persistent and mirroring across the asymmetric tasks. These patterns are similar to those observed in cats during locomotion with and without obstacles and reaching movements. These findings support the hypothesis that the profiles of muscle activations are qualitatively similar across quadrupeds during precise locomotor tasks.

**New and Noteworthy:** We characterize for the first time the spatiotemporal muscle activation in rat forelimb during precise asymmetric stepping on asymmetrically placed rungs. Similar to cats, the intralimb pattern of muscle activation in rats was stereotypical. The elements of this pattern were changing in a lateralized fashion based on the direction of the imposed asymmetry. The similarity of pattern to that of cats supports the idea of similarity of neural control across cat and rat species.

## Introduction

Locomotion is essential to the survival of all animals (Dickinson et al., 2000). From flagellar movement of unicellular organisms to the limbed locomotion generated by coordinated muscles of mammals, mobility offers distinct evolutionary advantages in finding prey, escaping predators, and courting mates. Yet, it is often challenging to navigate unpredictable terrain with obstacles and dangers requiring an animal to rapidly change direction or speed.

In vertebrates, the neural structures involved in locomotion are incorporated in a complex hierarchical system that seamlessly integrates commands from supraspinal motor pathways, intrinsic spinal rhythmogenic networks, and afferent sensory feedback (Grillner and Wallen, 1985; Yakovenko, 2011). At the top of motor hierarchy, the motor cortex has been shown to be an essential contributor for the accurate spatiotemporal control of discrete volitional limb movements, such as reaching for a target in both cats and primates (Kuypers, 1963; Lawrence and Kuypers, 1968; Lemon, 2008; Liddell and Phillips, 1944; Martin and Ghez, 1993, 1991), as well as for real-time modification of stereotypical rhythmic movements such as locomotion (Beloozerova and Sirota, 1988, 1993a; Drew, 1988, 1993; Drew et al., 2002, 2008b; Friel et al., 2007; Rossignol et al., 1996), and mastication (Hoffman and Luschei, 1980; Lund and Lamarre, 1974). In addition, single-unit recordings during such tasks have identified active pyramidal tract neurons (PTNs) that fire in specific patterns related to the change in task, such as during readjustment of reach trajectory (Yakovenko and Drew, 2015, 2009), when changing gait length or stepping over an unexpected obstacle (Beloozerova and Sirota, 1993a; Drew, 1993; Drew et al., 2008b, 2008a; Krouchev and Drew, 2013; Yakovenko and Drew, 2015), and during changes based on visual (Drew et al., 2008a) or proprioceptive feedback (Frigon and Rossignol, 2008). Moreover, these units fire discretely in relation to specific muscle groups during these tasks (Krouchev and Drew, 2013; Yakovenko and Drew, 2015) supporting the idea of control modules expressed within the nervous system.

The organization of downstream control elements has also been shown to express modularity, which was observed as the task specific grouping of muscles in stereotypical behaviors such as locomotion, scratching, and swimming (Bizzi et al., 2008, 1991; d’Avella et al., 2003; d’Avella and Bizzi, 2005; Tresch et al., 2002, 1999). Sequential muscle patterns are also observed in postural control of cats and humans (Ting and Macpherson, 2005; Yakovenko and Drew, 2009), in locomotion of cats and humans (Ivanenko et al., 2004; Lacquaniti et al., 2012), and in reaching movements of cats and primates, including humans (d’Avella et al., 2008, 2006; Drew et al., 2008b; Overduin et al., 2008; Yakovenko and Drew, 2015). The existence of the downstream modularity indicates that this organization principle has advantages for the integration of contributions from multiple neural pathways.

Yet, despite significant redundancy built into the motor control system, both central injuries and peripheral trauma can disrupt motor function. Cerebrovascular accidents or strokes are one of the primary causes of adult disability (Hodgson, 1998; Roger et al., 2011; Wasay et al., 2014), and can cause direct damage to the motor cortex. In hemiparetic patients this results in an asymmetric gait (Allen et al., 2011; Bowden et al., 2006; Brandstater et al., 1983; Wall and Turnbull, 1986), which may persist even after rehabilitation therapy (Ada et al., 2003; Patterson et al., 2010, 2008; Plummer et al., 2007). An interesting aspect of this hemiparetic gait is the heterogeneity within the patient population; either the paretic leg or the non-paretic leg may exhibit a shorter stride (Bowden et al., 2006; Wall and Turnbull, 1986). Step length asymmetry is an index that takes into account both the spatial (relative step lengths) as well as the temporal (relative foot-strike timing) aspects of the asymmetry in the hemiparetic gait with the lateralization of deficits (Malone and Bastian, 2010), and could perhaps serve to evaluate and design patient-specific therapy (Finley et al., 2015; Lauzière et al., 2014). Interestingly, some of the observed motor deficits in post-stroke patients can be explained by muscle synergy analyses, wherein their structure is disrupted by merging or fracturing of muscle synergies (Bowden et al., 2010; Cheung et al., 2012, 2009; Clark et al., 2010; Coscia et al., 2015). Further detailed characterization of the asymmetric locomotion in animal models could facilitate a more granular understanding of muscle coordination and motor modules in health and disease.

Here, we propose that rats are an eminently suitable model for such a multi-level approach. Firstly, rats are well-defined models of stroke and spinal cord injury (Farr and Whishaw, 2002). Secondly, they have a lisossomic cortical organization, which is appropriate for the use of modern electrode arrays (Whishaw et al., 2003). Finally, rats are also stereotaxically consistent animals, which makes them appropriate for the investigation of cortical and subcortical networks (Paxinos and Watson, 2007). However, rats are small quadrupedal animals that have not been neurophysiologically characterized to the same extent as cats or humans. There are clear similarities between the gait patterns of cats and rats, but there are also many differences. For example, rats walk with a crouched posture, bearing a greater proportion of their weight on the hind legs (Niederschuh et al., 2017). Rats exhibit freezing in the presence of predator odors (Griffith, 1920) and standing up driven by curiosity (Deschênes et al., 2012). There is a need to characterize locomotor behaviors and to establish equivalency or to investigate the disparity between cats and rats at the level of behavior and structure. This is the essential step toward establishing the rat as a model to further investigate the descending control of muscle coordination during normal and pathological conditions. Previously, we have used both extensive bilateral intramuscular implants and computational clustering analyses to document the sequence of events in the reaching movements and locomotion of cats (Yakovenko and Drew, 2015, 2009). We have also developed an instrumented pegway (Tuntevski et al., 2016) that permitted the testing of coordination during precise symmetric and asymmetric locomotor tasks in adult rats. Preliminary analysis of synergies were performed with the decomposition methods – PCA and NNMF – to study the methodological limitations and the persistence of synergies across symmetric and asymmetric locomotor tasks (Hanna and Yakovenko, 2021). The goal of this study was to determine the spatiotemporal progression of muscle activity during symmetric and asymmetric walking tasks in rats. We hypothesized that the sequence of muscle activations will be the same as those observed in cats.

## Methods

The experiments were carried out in eight adult female Sprague-Dawley rats (225-275g, 2-3 months old). Five animals had extensive bilateral implants and the remaining three had unilateral implants, allowing us a total of 13 implanted forelimbs for analysis. The animals were trained on the pegway task (described below) for one week prior to surgical implantation of EMGs. The surgical and experimental procedures followed the guidelines of and were approved by the WVU Institutional Animal Care and Use Committee (Protocol #15-0303).

### Surgery

All surgical procedures were performed under general anesthesia and under aseptic conditions. Anesthesia was induced by 5% isoflurane and maintained by 1-3% isoflurane with oxygen. A pre-emptive analgesic, Buprenorphine SR, was administered subcutaneously at 1.2 mg/kg, along with Dexamethasone at 2 mg/kg to reduce inflammation. For acute pain management and vasoconstriction, Lidocaine HCl at 4 mg/kg was injected at all incision sites prior to making incisions. The surface of the eyes was coated with petroleum jelly to prevent drying out. As described previously for mice (Pearson et al., 2005) and cats (Drew et al., 1986; Yakovenko, 2011; Yakovenko and Drew, 2009), forelimb muscles were implanted with pairs of Teflon-insulated, braided stainless steel wires (AS 633; Cooner Wire, Chatsworth, CA) for intramuscular electromyography (EMG). In each of the 13 limbs, the following representative muscles of the forelimb were chosen: *latissimus dorsi* (LtD) and *spinodeltoid* (SpD), which retract the limb as it is lifted in early swing; *cleidobrachialis* (ClB), which protracts the limb by flexing shoulder and elbow; *extensor carpi radialis longus* (ECR), which contributes to the transfer of the limb in swing with elbow flexion and wrist extension; *extensor digitorum communis* (EDC), which extends the wrist; palmaris longus (Pal), which generates wrist plantarflexion in the late swing preparing for the ground contact and the transition to stance; and *triceps brachii longus* (TrLo) with *triceps brachii lateralis* (TrLa) generating limb extension during limb placement and load-bearing in stance. These selected muscle actions are the same as those in our previous publication (Hanna and Yakovenko, 2021).

The rats were placed in a stereotaxic apparatus, and 4 stainless steel self-tapping anchor screws (FST, Foster City, CA) were placed at a depth of 0.25 mm in the cranium, approximately 3-4 mm anterior and 2-3 mm posterior to the lambda fissure, and 3-4 mm from the midline (note: the exact location of the screws was adjusted to accommodate individual animal morphology). Two 18-pin Omnetics connectors (A79038-001, MicroProbes, Gaithersburg MD) coming from the EMG implants were passed into custom 3D-printed housing, which was secured to the anchor screws with dental acrylic. This assembly of two connectors was attached to a single 32-channel interface board that directly plugged into a digitizing frontend (nano2stim+, Ripple, Salt Lake City, UT), allowing communication with the recording equipment (Grapevine Neural Interface Processor). Buprenorphine SR (1.2 mg/kg, SQ) was administered prior to the surgery and provided post-surgery analgesia for 72 hours. Ketoprofen (3 mg/kg) was administered as necessary to reduce any discomfort during the recovery period. Antibiotics were administered prophylactically for 5 days after surgery.

### Behavioral Task

Animals performed locomotor tasks on an adjustable 24-peg walkway instrumented with an array of force sensors embedded into pegs to record vertical forces from foot-strike (F_V_) see Figure 1 (Tuntevski et al., 2016). The position of pegs could be modified to impose either symmetric or asymmetric gait. We empirically determined 15 cm to be the preferred stride length for the rats by speeding up and slowing down locomotion on a treadmill until animals performed a continuous behavior without stopping or moving to the front of treadmill (Exer-3/6 Treadmill, Columbus Instruments, OH). Symmetric and asymmetric gait types were represented by three conditions shown in Figure 1B. The first symmetrical condition, S15 (black), corresponds to regular stride length symmetry with equal and out of phase stepping imposed by pegs set 7.5 cm apart, see Fig. 1B (middle). The second and third conditions impose asymmetric gait with preference assigned to either *ipsilateral* or *contralateral* limb by the asymmetric placement of pegs 9 and 6 cm apart (Fig.1B I9C6=red and I6C9=blue conditions) forcing asymmetric kinematic patterns. These stride conditions were alternated, and analyses were done relative to the EMG implantation side. Animals were rewarded for the successful completion of each trial with verbal encouragement and food (Frootloops, Kellog, Battle Creek, MI).

### Data Collection

After a week of recovery, implanted animals performed the locomotor tasks while tethered to the acquisition system (Grapevine Neural Interface Processor, Ripple, Salt Lake City, UT). EMGs and ground reaction forces were recorded simultaneously. Single ended recordings from the implanted pairs of electrodes were conditioned and sampled at 30 kHz using a digitizing frontend attached to the connector on animal’s head. The synchronized ground reaction forces generated by contacts of both forelimbs and hindlimbs with each peg were conditioned and sampled at 1 kHz with an analog-to-digital card frontend. The dataset included only consecutive steps where the animal walked continuously without pausing or running.

### Signal Processing

Digitized vertical force (F_v_) recorded with an array of force-sensing resistors were analyzed using a custom script in MATLAB (MathWorks, Natick, MA). The time of stance onset (red triangles) for forelimbs was semi-automatically marked using a supervised onset detection method (Yakovenko et al., 2005) for all steps in each trial. Data from inconsistent steps when subject stopped every 2 steps (less than 3 steps in a bout) or slowed down or sped up for more than two standard deviations from the average step period. The initiation of the swing phase can be clearly seen from the deflection in the force profile. These events were marked manually as ‘offsets’ (blue triangles) in the custom interactive analysis environment. The complete stance offset was manually marked with a ‘peak’ marker (green triangles) (Figure 2).

The digitized EMG data were first filtered with a high-pass second order Butterworth filter with a cutoff frequency of 20 Hz, standardized by subtraction of mean value, and then subjected to a low-pass second order Butterworth filter with a cutoff frequency of 100 Hz, using custom MATLAB® (MathWorks, Natick, MA) scripts. Each EMG signal was normalized to the peak-to-peak value of the corresponding symmetrical walking condition.

### Intralimb Coordination

We analyzed ipsilateral muscle burst latencies during swing phase in about 80 steps per limb. The EMG signals were aligned with the onset of ipsilateral limb transfer and normalized to the duration of the ipsilateral swing phase. The values of muscle burst latency are represented as a percentage of swing phase where 0% is the start and 100% is the end of swing. Additionally, 10% padding was added before onset of limb transfer and after stance onset for visualization. Muscle burst latencies were manually identified and measured on each limb EMG signal profile (Figure 3A, arrows). On average, eleven measurements per muscle were included in the analysis. Signals were excluded if there was evident electrode failure. The values of muscle burst latency were analyzed with a one-way ANOVA.

Changes in EMG and ground reaction force (GRF) amplitudes caused by an imposed asymmetry were analyzed using a binned method. Signals were averaged across about 80 steps per limb, spatiotemporally normalized and divided into 10 bins of equal duration. Normalization of signals was done (1) temporally of the full step cycle starting with swing onset and (2) to the peak-to-peak values of the corresponding signal in the symmetric condition. Correspondingly, Bin 1 represents the beginning of swing phase while Bin 10 represents the end of stance phase. The average normalized amplitude of the portion of the signal captured in each bin was calculated, and these values are represented with box plots for each condition (n=13) (**Figure 5**). A descriptive post-hoc analysis using one-way ANOVA was performed per bin to test for amplitude differences across conditions. Since this is a descriptive post hoc analysis, a correction for multiple comparisons was not used.

### Interlimb Coordination

The asymmetry of locomotion was further described with an interlimb analysis of the double stance phase, or the period when both limbs are in stance. Each step cycle contains two double stance phases which can be described as either (1) leading or (2) trailing. Leading double stance starts with ipsilateral stance onset and ends with contralateral stance offset, while trailing double stance starts with contralateral stance onset and ends with ipsilateral stance offset (see Figure 2). Although intralimb analyses in this study were performed in each limb, interlimb analysis was done per-animal to prevent repeating data (n=8).

Double stance times were calculated using the ground reaction force ‘onset’ and ‘peak’ events in both limbs (see above *Signal Processing*). Average leading and trailing double stance times for about 80 steps in each animal and each condition are compared in **Figure 6**. Leading and trailing double stance times are compared within each condition using a two-tailed t-test to determine a difference within each condition. Double stance time differences, calculated as trailing minus leading in each condition, are compared across conditions using a one-way ANOVA. Additionally, the double stance time difference for the symmetric condition is compared to a mean of zero using a two-tailed t-test, and the absolute values of the double stance time differences in the asymmetric conditions were compared using a two tailed t-test. The level of significance was set at 5% for all tests, and the differences are shown with asterisks (*).

## Results

Both temporal and amplitude analyses of forelimb muscles relative to the vertical ground reaction forces were performed in this study.

### EMG Temporal Profile

The muscle activity profile follows a standard sequence indicated by the increasing onset latency (Figure 3). Limb retractors, *LtD* (mean ± SD values are 1.4 ± 1.4%) and *SpD* (6.1 ± 5.9%), are the first to activate and lift the limb off the ground. Then, limb protractors, represented by *ClB* (23.1 ± 12.7%) and *ECR* (26.9 ± 9.1%) transport the limb in swing. Then, wrist extensors represented by *EDC* (48.9 ± 8.9%) help with toe clearance over ground. Finally, wrist plantar flexor *Pal* (60.5 ± 7.4%) and elbow extensors *TrLo* (64.1 ± 9.7%) and *TrLa* (81.5 ± 12.1%) prepare for foot contact and limb loading in stance (Figure 3).

The temporal sequence of muscle activations is similar during both symmetric and asymmetric conditions. Both ipsilateral and contralateral limb preference conditions correspond to similar spatiotemporal pattern of muscle activations. *LtD* (0.4 ± 3.9%, 0.8 ± 2.7%) and *SpD* (5.4 ± 4.0%, 5.1 ± 5.3%) are activated first, followed by limb protractors *ClB* (23.5 ± 12.8%, 21.3 ± 11.0%) and *ECR* (27.8 ± 7.4%, 28.9 ± 8.5%). Then, wrist extensor and dorsiflexor *EDC* (49.1 ± 9.7%, 49.5 ± 9.1%) is activated, followed by wrist plantar flexor, *Pal* (59.6 ± 6.2%, 61.6 ± 5.8%), and elbow extensors *TrLo* (67.2 ± 6.9%, 64.4 ± 8.8%) and *TrLa* (71.3 ± 30.3%, 76.0 ± 15.3%) (mean ± SD values indicated for ipsilateral and contralateral preference conditions, respectively) (Figure 4).

Muscles can be separated into four distinct groups that represent the lift, flexion and transport, toe clearance, and prepare-for-contact stages of swing phase (Figures 3B and 4B). This grouping that is maintained across symmetric and asymmetric walking supports the hypothesis that the same muscle synergies are used to control symmetric and asymmetric locomotion.

### EMG Amplitude and Temporal Analysis

As expected, the differences in the asymmetric conditions are generally opposite (mirroring) relative to the symmetric condition. Normalization masks these differences in the ipsilateral vertical force (*iF*), and instead they are evident in the contralateral vertical force (*cF*) bins 1,5-8. A forward temporal shift in contralateral swing onset in the I9C6 (red) condition is represented by a decreased vertical force amplitude of the limb with respect to the symmetric condition (*cF;* p < 0.001, <0.001, Bins 6 and 7, respectively). A mirrored, delayed temporal shift in contralateral swing onset in the I6C9 (blue) condition is represented by increased vertical force with respect to the symmetric condition (*cF;* p < 0.001, 0.001, Bins 6 and 7, respectively). In contralateral stance onset, stance offset, and swing onset, the forward temporal shift of the I9C6 step cycle and delayed temporal shift of I6C9 step cycle is maintained (*cF*; p <0.05, <0.001, <0.001 Bins 1, 5 and 8, respectively). This confirms consistent and opposite changes in the asymmetric conditions promoting asymmetric loading of limbs.

The ipsilateral EMG activity shifted in the asymmetric conditions. During late ipsilateral swing, there was decreased amplitude in one of the limb retractors (SpD, <0.001, Bin 1), increased amplitude of shoulder and elbow flexion in the I9C6 (red) condition (*ClB*; p <0.001, <0.001, <0.01, Bins 3-5, respectively). The pattern indicates faster shoulder and elbow flexion during limb transport with respect to the opposite condition, and, thus, faster ipsilateral stance onset in this condition. At the onset of stance, the amplitude of plantarflexors (*Pal*) and limb extensors (*TrLo*) increased (*Pal*; p = 0.062, 0.007, 0.007 in Bins 5-7, respectively and *TrLo*; p = <0.001, <0.001, <0.01 in Bins 5-7, respectively). This indicates that the ipsilateral stance was accelerated in this condition promoting the loading of the ipsilateral limb.

Oppositely in the I6C9 (blue) condition, the amplitude decreased in shoulder and elbow flexors (*ClB*; p <0.001, <0.001, <0.01, in Bins 3-5), plantarflexors (*Pal*; p = 0.062, 0.007, 0.007 in Bins 5-7, respectively), and limb extensors (*TrLo*; p = <0.001, <0.001, <0.01 in Bins 5-7, respectively) indicating a delay in the transition to ipsilateral stance and the prolongation of contralateral support phase. These changes relative to the symmetrical condition are opposite of those seen in I9C6. Thus, the two kinematic conditions constitute simulated limb preference.

### Double Stance Analysis

The comparison of double stance phases and their asymmetries in different conditions are shown in **Figure 6**. Double stance phases of leading and trailing limbs are, as expected, not different in the symmetric condition (Fig. 6A, black, p = 0.282). These phases are different in the symmetric conditions (Fig. 6A, p <0.001, <0.001 for red I9C6 and blue I6C9 respectively). The difference between the trailing and leading phases in Fig. 6B was consistent between three conditions (pair-wise comparisons between symmetrical and two asymmetric conditions: p<0.001 for I6C9 and <0.001 for I9C6). For example, in the red condition (I9C6), the ipsilateral limb initiates swing soon after the trailing limb initiates stance. This configuration of limbs between the neighboring pegs is less challenging than the one when limbs are placed on pegs that are far from each other in the trailing double-stance phase. The ipsilateral trailing double stance duration is, thus, longer to accommodate the increased need for support.

## Discussion

In this study, we document for the first time a stereotypical pattern of muscle activations during symmetric and asymmetric precise stepping in rats. Previously, the pattern of sequential muscle activation has been described in cats during the symmetrical locomotion on a treadmill with and without obstacles and during the reaching to a lever movement (Forssberg et al., 1980; Yakovenko et al., 2011). While kinematic analysis of stepping in rodents has been extensively based on motion tracking and even exoskeleton use (Garrick et al., 2021; Hewitt et al., 2018; Song and Hogan, 2015), the description of muscle coordination in the context of the manipulation of stepping symmetry is a novel result in this study. A stereotyped spatiotemporal sequence of activation in representative forelimb muscles was persistent and mirroring across the asymmetric tasks.

### Muscle groups for symmetric and asymmetric gait

Similar to cats, rats use the same muscle groups or synergies to execute swing phase of locomotion (Fig. 3&4), which can be further subdivided into *limb lift*, *flexion*, *transport*, and *preparation for stance*. The muscle recruitment onsets are organized in the following sequence: limb retractors (*LtD, SpD*), elbow flexors (*ClB, ECR*), wrist dorsi-/plantar- flexors (*EDC, Pal*), followed by load bearing muscles during stance (*TriLo, TriLa*). The burst duration of *ClB* and *ECR* were similar, unlike in cats, and likely associated with the requirement to maintain flexion for ground clearance in the crouched posture of rats. While cortical activity during the precise stepping in rats has not been reported, a sequential pattern of cortical discharge has been observed in cats and was associated with the coordination of muscles during reaching movements (Yakovenko et al., 2011) and also during locomotion (Yakovenko and Drew, 2015). It is likely that the conservation of the spatiotemporal activation of muscles is likely accompanied by the similar cortical dynamics in these two species.

While multiple feedforward and feedback as well as spinal and supraspinal pathways are converging to provide online control of stepping and contributing to muscle activations (Rossignol, 1996), the reason for the expression of integrated patterns at the cortical level is unknown. It is curious to note that the postural control in decerebrated cats is compromised despite the fact that their sensory pathways are intact; thus, the contribution of the descending control inputs are essential. Previously, the existence of these persistent behavioral templates at multiple levels in the control hierarchy was attributed to the challenging task of integrating limb and whole-body posture control (Yakovenko and Drew, 2015). This hypothesis is supported by the view of internal representations of limb and body dynamics necessary for the planning and execution of movement in the presence of transmission delays and unpredictable interactions with the environment (McNamee and Wolpert, 2019; Wolpert et al., 1995). While this holistic view on control suggests complex representation of the whole system at the high levels in the hierarchy, it includes the additive layered organization of multiple feedback loops. The output of such a system is an emergent property of the interactions between descending and ascending pathways.

### Low-dimensional and high-dimensional representation of control

The existence of spatiotemporal templates composed of the temporal sequences of muscle group activity that can be independently modified in task-dependent manner contradicts the idea of the general low-dimensional representation of control. The support for this alternative typically comes from the studies using statistical dimensionality reduction techniques in reaching, postural, and locomotor tasks (Churchland et al., 2012; Xing et al., 2019). Almost any pattern can be mathematically expressed as a low dimensional signal using, for example, a popular dimensionality reduction method like principal component analysis (PCA) (Patla, 1985; Tresch et al., 2006; Tresch and Jarc, 2009). One example of compact control is the use of limb speed to control turning during locomotion. Using a relatively simple interlimb speed difference in models of pattern generation, we can express a change in the heading direction for turning (Yakovenko et al., 2018). The simple command expressing limb speed, a low-dimensional input to the spinal synergies, would then be transformed through the CPG network into a high-dimensional spatiotemporal spinal output defined by the number of controlled muscles (Desrochers et al., 2019). Another example of a low-dimensional representation is the postural and movement control using simplified “anchor behaviors”, for example, spring-loaded inverted pendulum, which can be embedded in predictive and reactive neural computations (Full and Koditschek, 1999). This low-dimensional control can be consequently evident in the decomposition of muscle activity, and it can be functionally demonstrated in the ability of animals to walk with minimal cortical contribution. Yet, these observations do not negate the presence of high-dimensional signals in the descending pathways. The variable cortical contribution during locomotion has been shown by the tuning of cortical neurons to the phasic muscle activity (Beloozerova and Sirota, 1993b; Drew, 1988). In this context, the cortical contribution was essential in the tasks requiring precision and dexterity (Georgopoulos and Grillner, 1989), for example, walking on ladders, which is similar to the task in this paper, that can be disrupted by the transection of the corticospinal tract.

The existence of low-dimensional representations, like the CPG, does not necessarily imply that the neural system tends to generalize or use exclusively low-dimensional computations. In tasks where dexterous manipulations may be required the associated high-dimensional control may co-exists to expand the repertoire. For example, the general locomotor behavior can be combined with precise stepping over obstacles that requires detailed representation of force generation responsible for the subphases of swing phase or reaching movements (Yakovenko et al., 2011; Yakovenko and Drew, 2009). The result in this study supports the existence of spatiotemporal templates that allow these dexterous movements based on the similarity of muscle grouping across species. While these templates can be expressed through a low-dimensional structure, it is likely that the control pathways responsible for precise limb placement during symmetric walking are also responsible for precise limb placement during asymmetric conditions by modifying the same motor program. However, the relative contribution of different mechanisms may change depending on the dexterity of motor task. These analyses support the idea that both animal models use the same temporal pattern of muscle activity as seen previously in cats during locomotion (Drew, 1993; Drew et al., 2008a; Lavoie and Drew, 2002; Yakovenko and Drew, 2015) and reaching (Holdefer and Miller, 2002; Yakovenko et al., 2011; Yakovenko and Drew, 2015). Recent synergy analysis with the dimensionality reduction techniques, PCA and non-negative matrix factorization (Hanna and Yakovenko, 2021), have also supported the existence of a consistent spatiotemporal activity template in this task, which is similar to that of cats. Overall, these observations support the similarities in motor coordination across rat and cat animal models.

## Conclusions

This study is first to our knowledge to document the forelimb muscle activity during a precise stepping task in rats. Moreover, the pattern is similar to that observed in cats during both reaching movements and locomotion with obstacles. The similar organization of the motor activity across cat and rat species supports the similarity of neural control and translational value of research in both models.

---

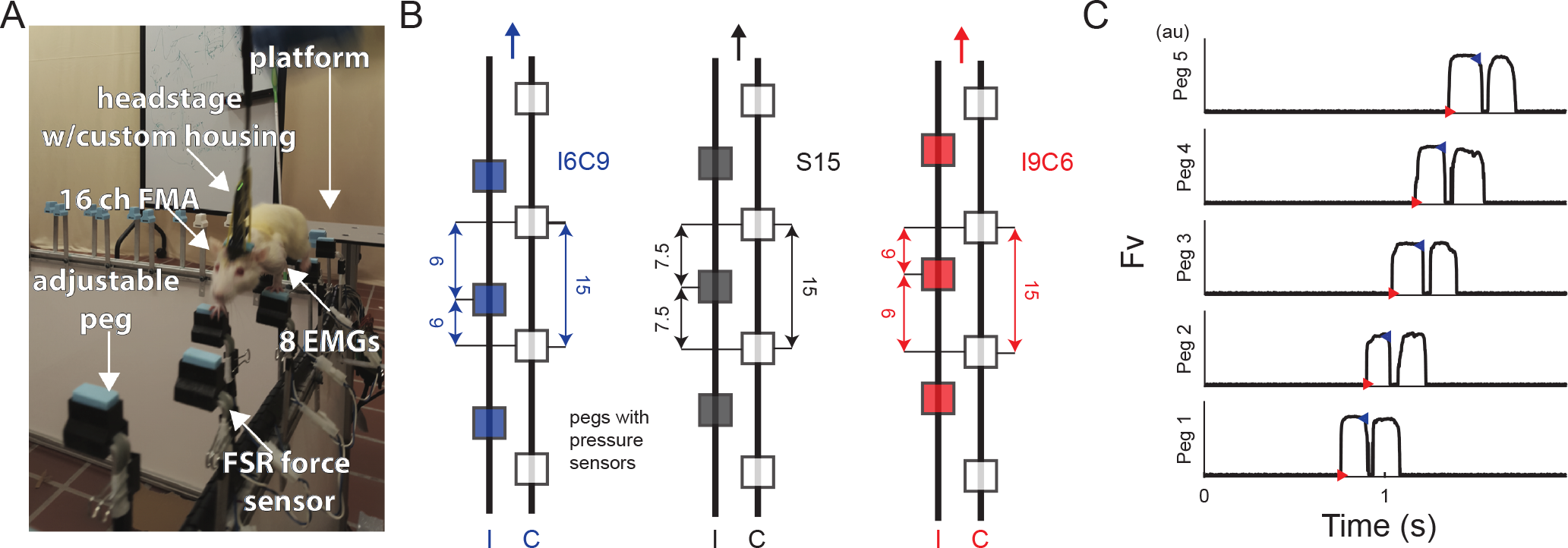

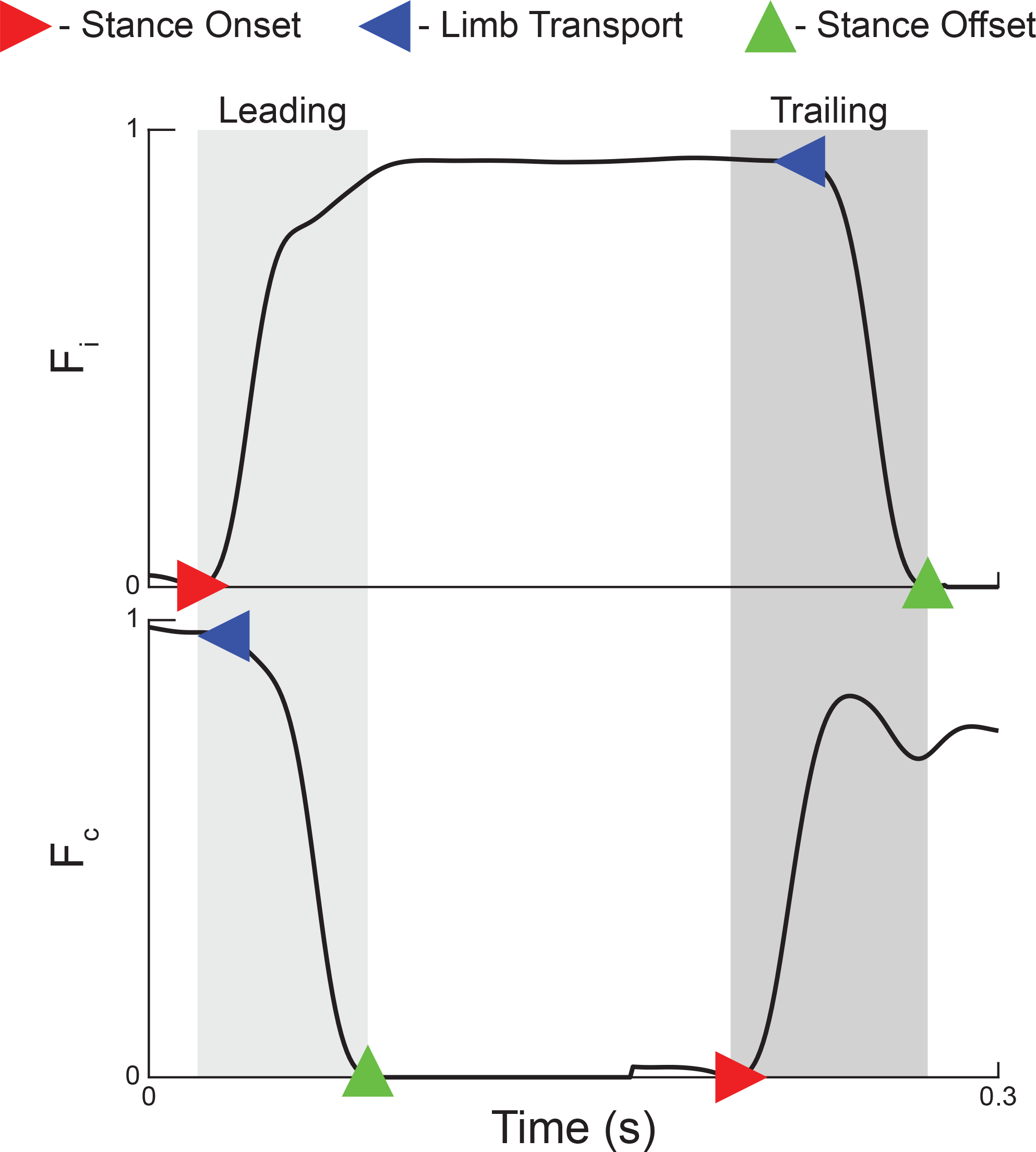

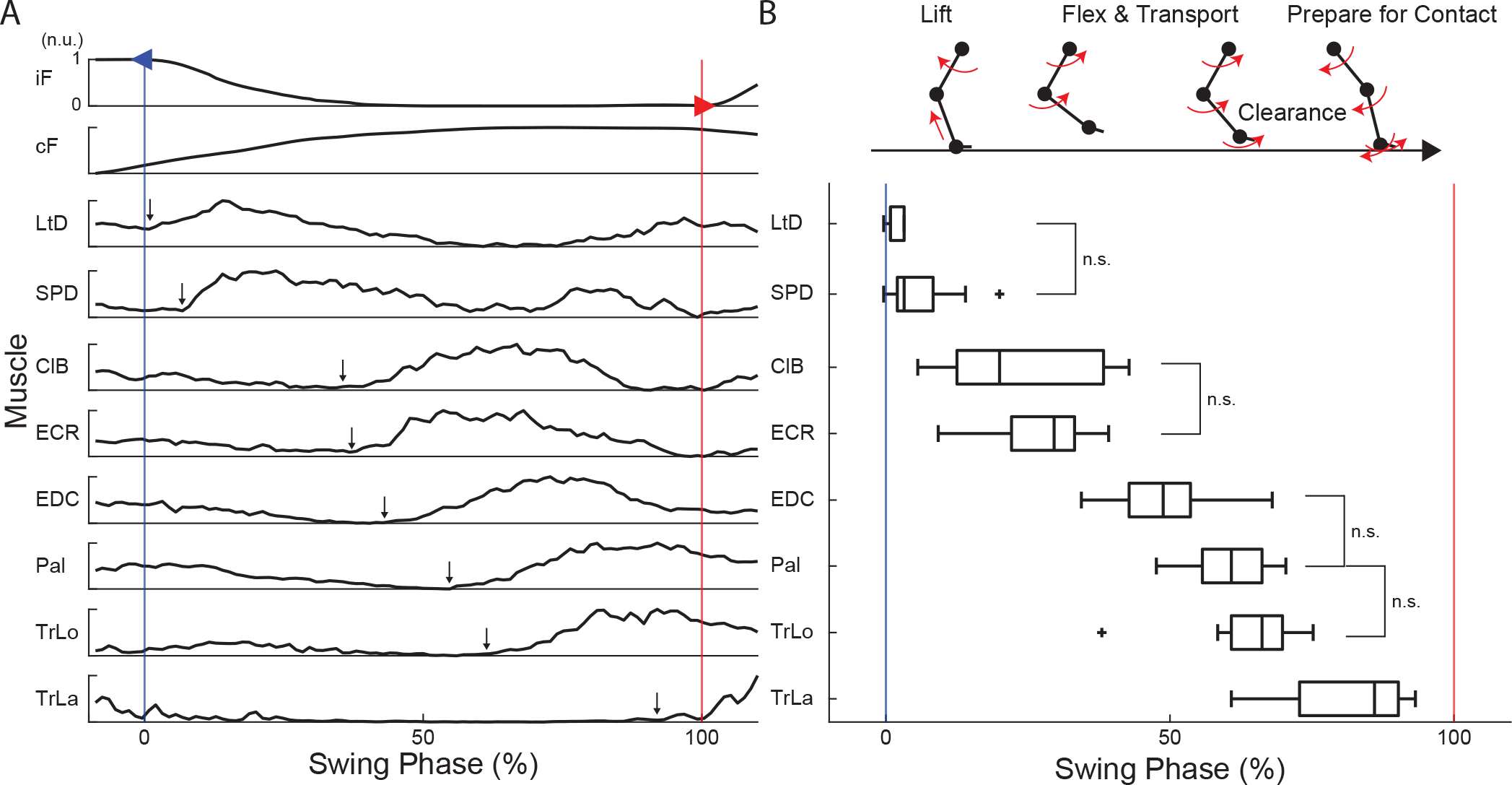

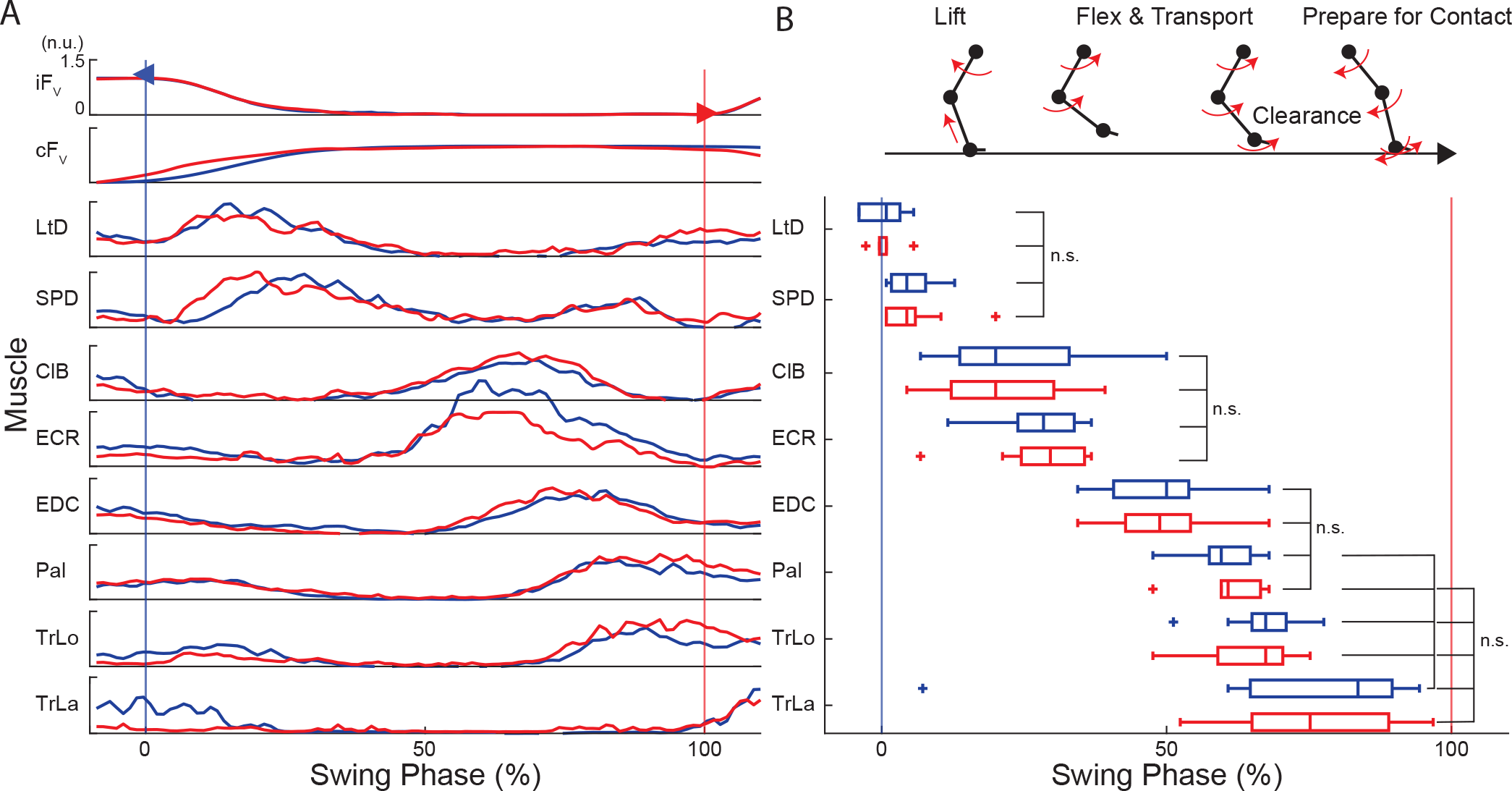

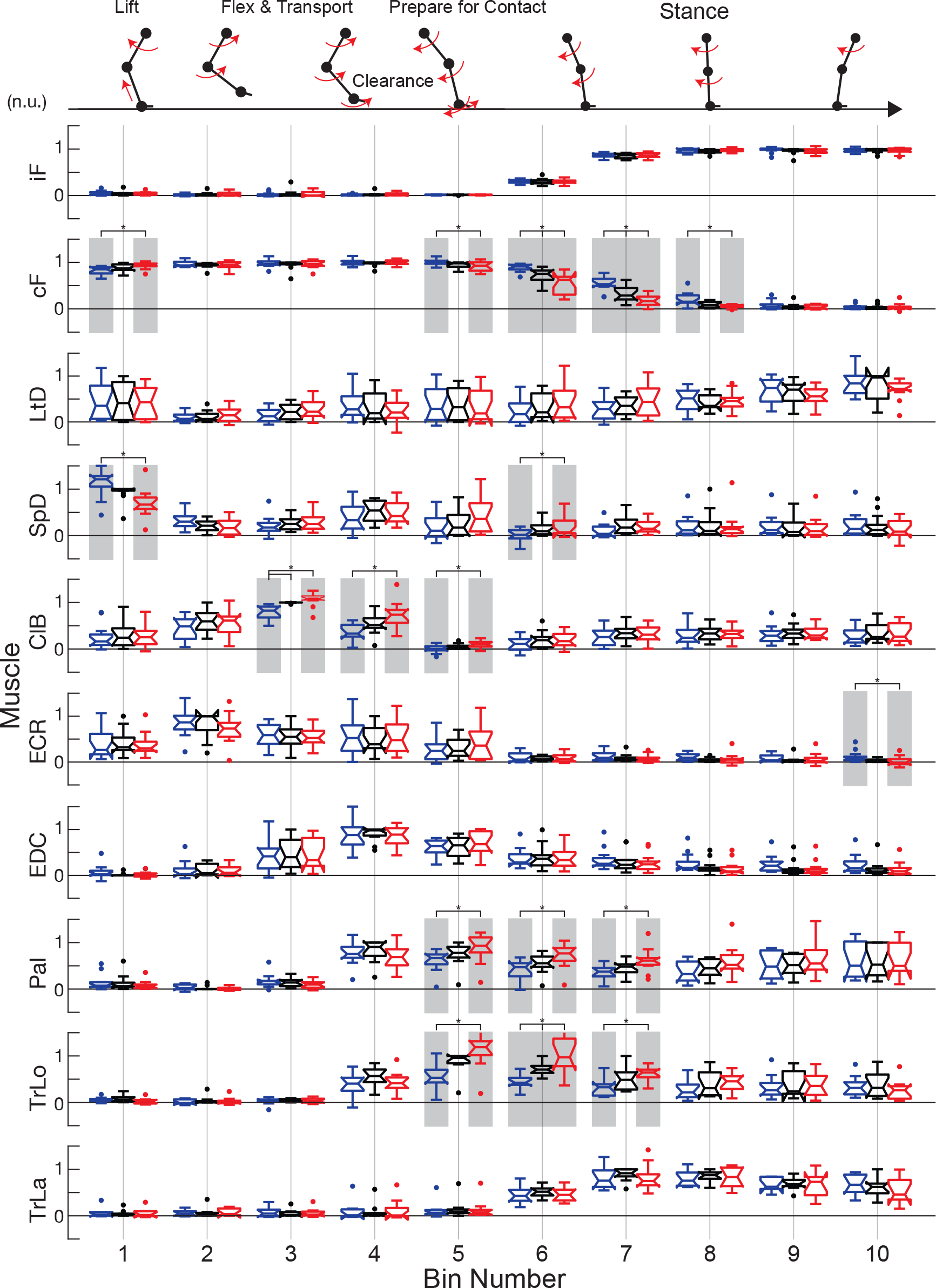

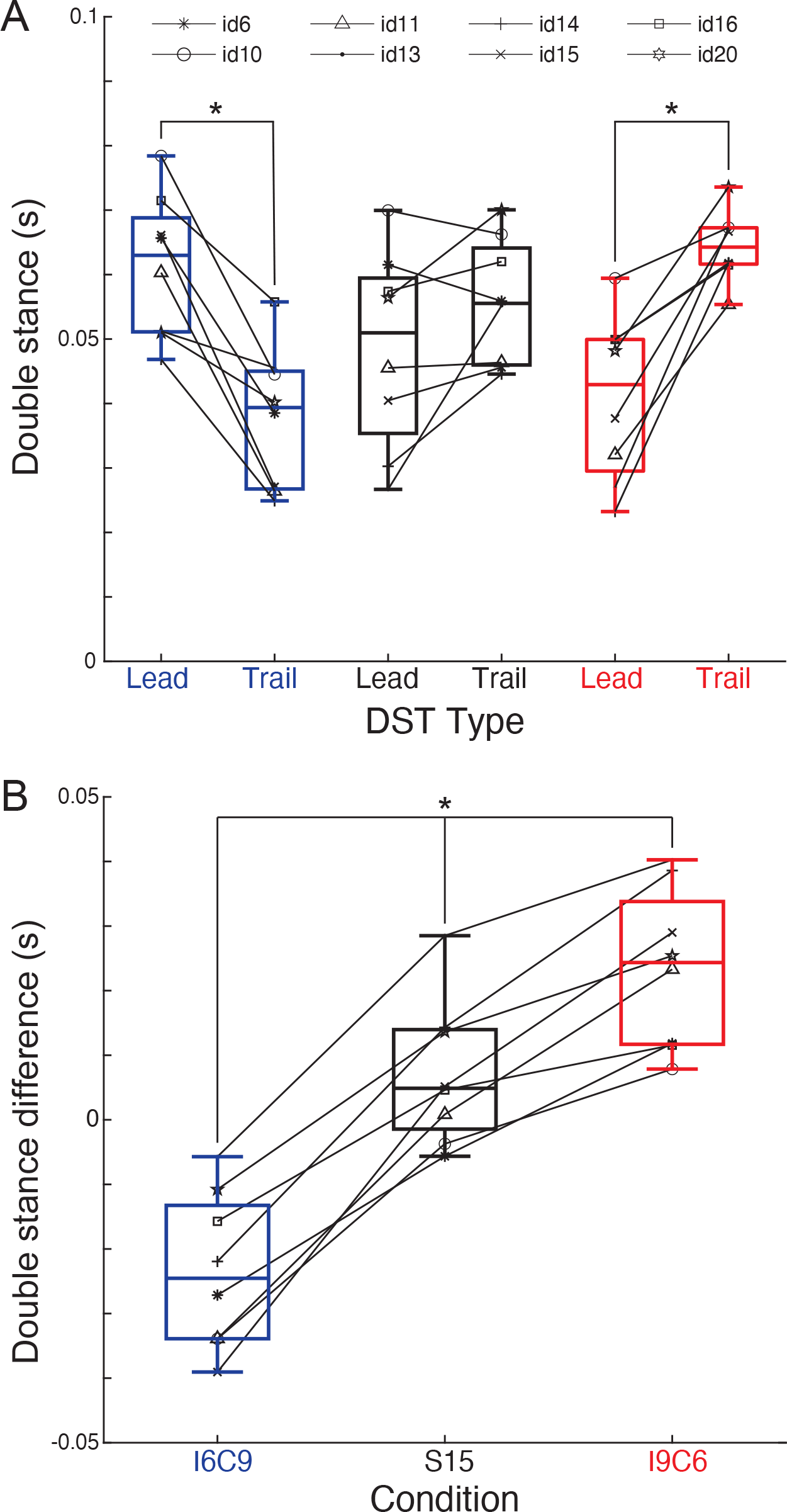

